# Cellular exocytosis gene (EXOC6/6B): a potential molecular link for the susceptibility and mortality of COVID-19 in diabetic patients

**DOI:** 10.1101/2020.06.25.170936

**Authors:** Mahmood Yaseen Hachim, Ibrahim Yaseen Hachim, Saba Al Heialy, Jalal Taneera, Nabil sulaiman

## Abstract

Diabetes is one of the most critical comorbidities linked to an increased risk of severe complications in the current coronavirus disease 2019 (COVID-19) pandemic. A better molecular understanding of COVID-19 in people with type diabetes mellitus (T2D) is mandatory, especially in countries with a high rate of T2D, such as the United Arab Emirates (UAE). Identification of the cellular and molecular mechanisms that make T2D patients prone to aggressive course of the disease can help in the discovery of novel biomarkers and therapeutic targets to improve our response to the disease pandemic. Herein, we employed a system genetics approach to explore potential genomic, transcriptomic alterations in genes specific to lung and pancreas tissues, affected by SARS-CoV-2 infection, and study their association with susceptibility to T2D in Emirati patients. Our results identified the Exocyst complex component, 6 (*EXOC6/6B*) gene (a component for docks insulin granules to the plasma membrane) with documented INDEL in 3 of 4 whole genome sequenced Emirati diabetic patients. Publically available transcriptomic data showed that lung infected with SARS-CoV-2 showed significantly lower expression of *EXOC6/6B* compared to healthy lungs.

In conclusion, our data suggest that *EXOC6/6B* might be an important molecular link between dysfunctional pancreatic islets and ciliated lung epithelium that makes diabetic patients more susceptible to severe SARS-COV-2 complication.

## Introduction

Since its appearance in late 2019, the COVID-19 outbreak represents a significant public health crisis with a heterogeneous impact on different population subgroups[1-3]. Indeed, while most of the patients show only a mild-moderate form of the disease, around 10-15% of patients suffer from a more aggressive form of the disease presented as severe to critical symptoms[4]. Many reports highlight that patients with specific comorbidities are more susceptible to the critical form of the disease compared to the general population [5]. Therefore, further investigations and early identification of those subgroups might be essential to improve the outcome of these vulnerable patients.

Indeed, T2D, in addition to hypertension, coronary heart diseases, and obesity is considered as the most critical risk factors of the severe form of SARS-CoV-2 infection and is associated with higher mortality [6, 7]. Interestingly, while several reports showed a similar prevalence of COVID-19 infection in diabetic patients compared to the general population, those patients were more vulnerable to the adverse outcomes with high mortality rates [8, 9]. It is not apparent whether T2D independently contributes to this increased risk or whether it is the combination of T2D with other risk factors such as cardiovascular disease (CVD), obesity, and hypertension[10]. A few reports documented that COVID-19 patients with diabetes showed specifically severe pneumonia, the release of tissue injury-related enzymes, excessive uncontrolled inflammation responses and hypercoagulable state associated with dysregulation of glucose metabolism, rendering patients with diabetes more susceptible to an inflammatory storm which eventually may lead to rapid deterioration of COVID-19[11].

Additionally, it has been reported that almost 50% of COVID-19 patients in Wuhan suffer from transient hyperglycemia, which could be due to SARS-CoV-19 binding to host receptors in pancreatic islets [2]. Moreover, many diabetic patients with COVID-19 had uncontrolled hyperglycemic states even when blood glucose management strategies were applied [12]. Such uncontrolled hyperglycemic levels will expose those patients to secondary infections along with higher mortality risks.

Due to the high prevalence of T2Din the U.A.E. (the 10th highest diabetes prevalence globally) [13] and the limited comprehensive research to decipher the pathophysiological mechanisms that govern the relationship between diabetes and COVID-19 [14], additional efforts is needed to investigate the association of clinical course between diabetes and COVID-19. Moreover, identification of the cellular and molecular mechanisms that render T2D patients prone to an aggressive course of the COVID-19 is an urgent matter for a better understanding of the COVID-19 pathogenesis.

Given the common embryonic developmental origin of the lung and pancreas, it is not surprising that common mechanisms driving COVID-19 infection might be analogous in the two organs, which lead to the deterioration of COVID-19 diabetic patients. Therefore, we employed a system genetics approach to explore potential genomic, transcriptomic alterations in genes specific to lung and pancreas tissues affected by COVID-19 infection, and are associated with susceptibility to T2D in Emirati patients. Our results identified *EXOC6/6B* gene (a component of the exocyst complex that docks insulin granules to the plasma membrane) with documented SNPs associated with diabetes in Emirati patients. Publically available transcriptomic data showed that lung infected with SARS-COV-2 showed significantly lower expression of EXOC6 compared to healthy lungs.

## Methodology

### Study Design, Population, and Settings

The details of the study design and sampling of the UAEDIAB study are described elsewhere [15]. The study was approved by the Ethics Committee of Sharjah University and the Ministry of Health Research Ethics Committee (R.E.C. number MOHAP/DXB/SUBC/No.14/2017). All participants had consented to the usage of their collected data and samples.

### Impact of SARS-COV-2 infection on genes involved in insulin secretion in hPSC-derived pancreatic organoid derange

To confirm the link between T2D and COVID-19, a transcriptomics dataset (GSE151803) of SARS-CoV-2 infected hPSC-derived pancreatic organoids[16] were retrieved from Geo Omnibus dataset https://www.ncbi.nlm.nih.gov/geo/. The R.N.A. sequence expression was explored through the online tool (BioJubies) to compare the differentially expressed genes and identify top pathways enriched in SARS-CoV-2 infected versus mock-infected samples.

### Targeted Next-Generation Sequencing from T2D Emirati patients

Four Emirati diabetic patients (age 60 ± 1, HbA1c >10.5%), fasting blood glucose values >13 mmol/L, BMI >40 and weight 120 ± 3 kgs) were subjected for targeted DNA next-generation sequencing using the S5 semi-conductor based DNA sequencer as described previously [26][17]. Briefly, the targeted coding-exome sequencing approach was designed multiplex primers that map across key genes along the coding region of the genome. The two patients were multiplexed on one Ion Chip. A pooled barcoded amplicon-tagged library generated using Fluidigm Access Array (Fluidigm Europe B.V, Amsterdam, The Netherlands) was diluted and subjected to emulsion polymerase chain reaction (PCR) with Ion SphereTM particles with Ion Template OT2 200 Kit (Ion OneTouchTM system) following the manufacturer’s instructions (Thermo Fisher, MA, U.S.A.). The pooled samples were sequenced using 540 Ion Chip with Ion 200 Sequencing kit on the S5 sequencer following manufacturer instructions (Thermo Fisher). The total mapped reads obtained were around 40 million per patient, with 109 mean depth and 94% uniformity coverage. 92.4% of the amplicons were aligned to the reference genome (HG19 build). The AQ20 quality mean length was 138bp, and the mean raw accuracy was 98.6%. The mean coverage across the amplicon was around x130. The bioinformatics analysis was carried out by first aligning the data using the B.W.A. alignment algorithm, followed by sequence filtering using SAMtools. Mutations file (V.C.F.) was generated using vcftools and Integrated Genome Viewer.

### RNA-sequence expression from human pancreatic islets

RNA-sequence expression data from human pancreatic islets were retrieved from a publicly available database (https://www.ncbi.nlm.nih.gov/geo) (GEO, accession number: GSE50398). As previously described[18], pancreatic islets were isolated from 89 cadaver donors; of them, 67 were nondiabetic and 12 were patients with T2D.

### Identification of common genes expressed by lung and pancreas

Publicly available tissue gene expression dataset (GTEx) was explored by BioJubies to compare the differential expressed genes of 427 healthy lung samples with 248 healthy pancreases (Figure 1). Genes with less than 2-fold changes (with adjusted p-value more than 0.05) were considered not specific to the lung or pancreas and thereby selected as shared genes between the two tissues.

**Figure 1:**
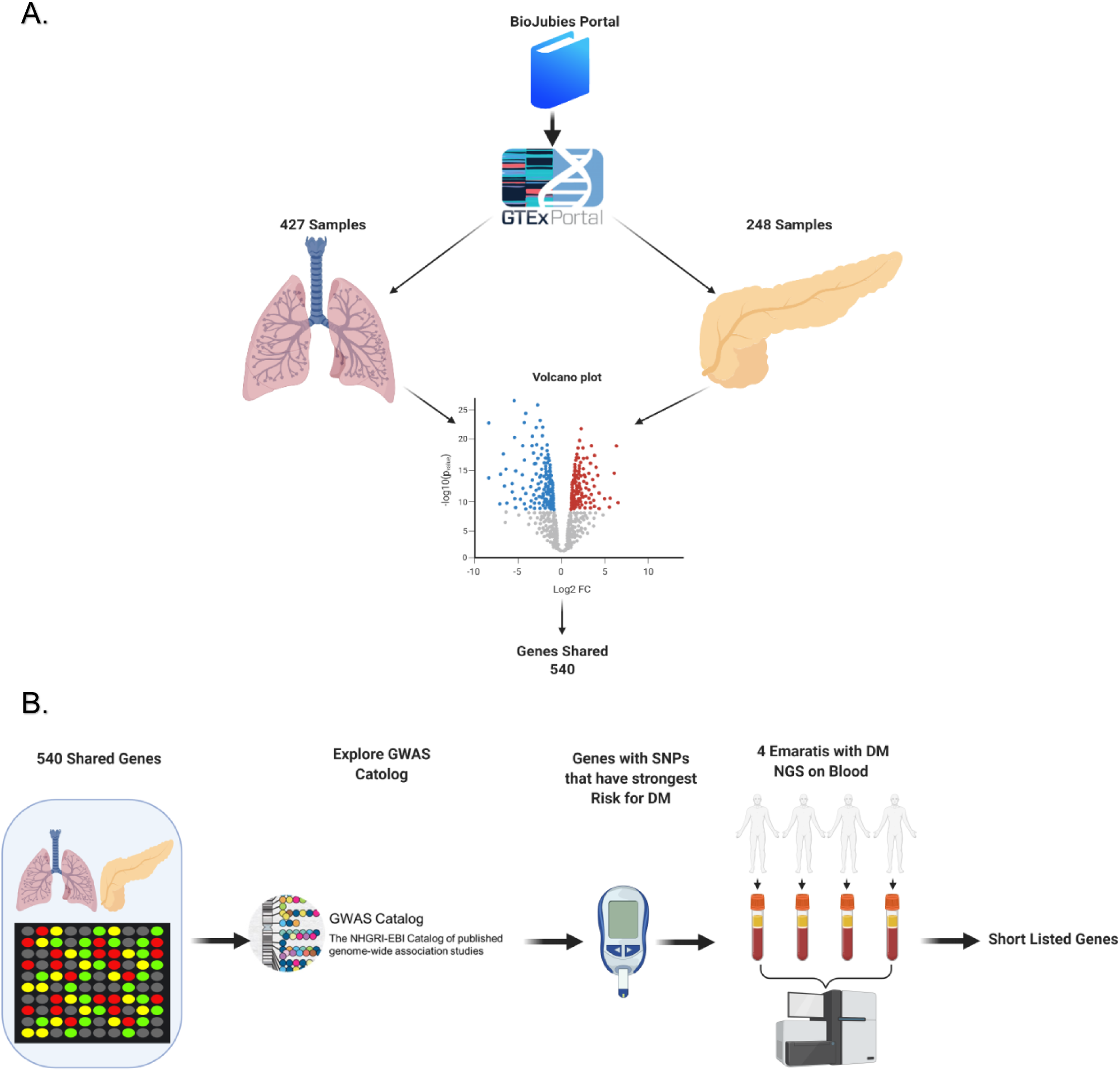
Flowchart for the identification of shared genes in lung and pancreas genes (A) and the association between SNPs and risk of T2D in Emirati diabetic patients (B).

### Identification of genes with T2D associated-SNPs in Emirati diabetic patients

The shortlisted of shared genes to lung and pancreas tissues were screened using genome-wide association studies (GWAS) database [19] to identify genes with T2D associated-SNPs. The final list of genes containing SNPs was further explored in the genomic profiling of the target next-generation sequence obtained from T2D Emirati patients (Figure 2).

**Figure 2:**
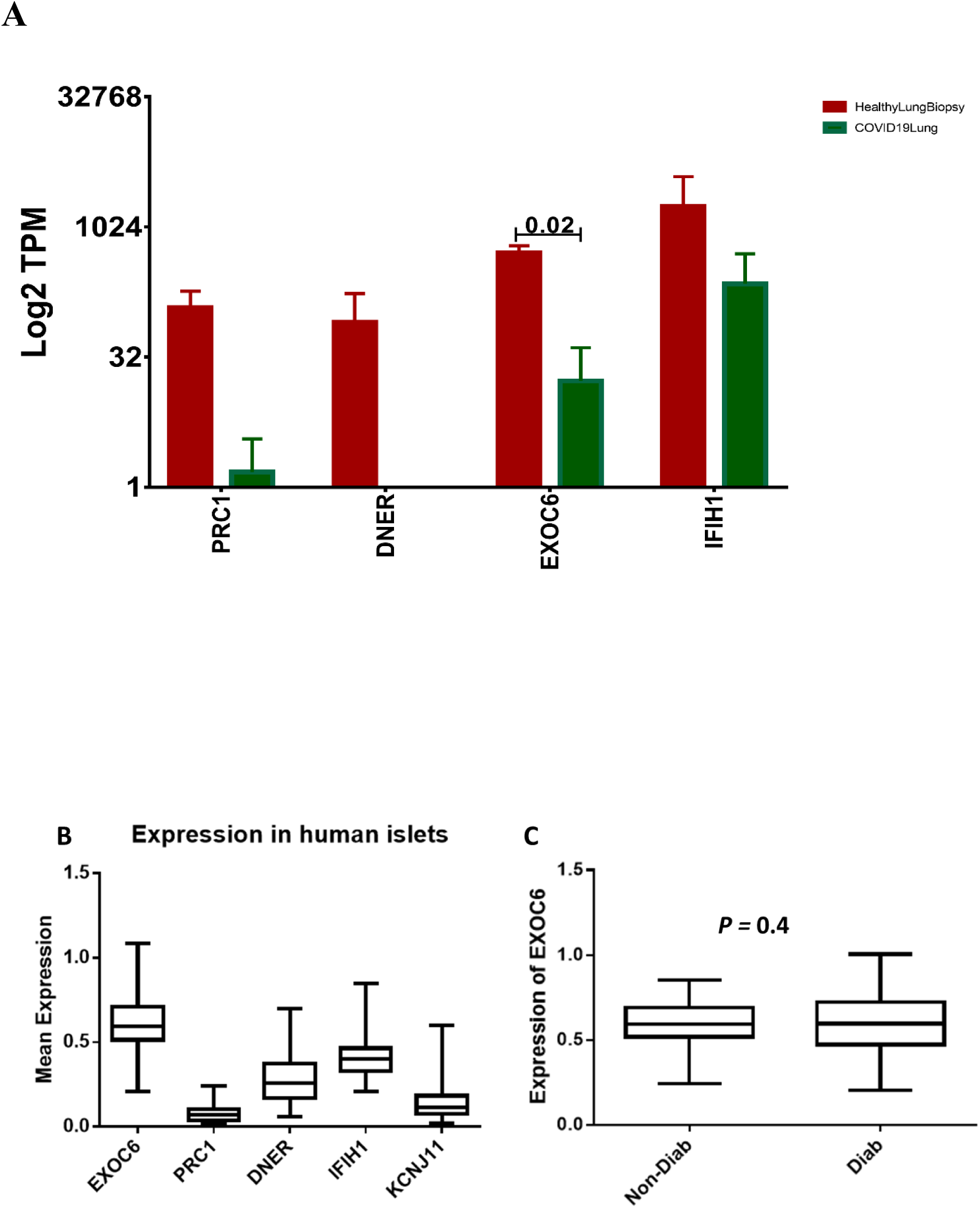
Expression of **of PRC1, DNER, *EXOC6/6B*, and IFIH1 in lung tissues infected with of SARS-COV-2 or human pancreatic islets.** **(A)** mRNA expression analysis of *PRC1, DNER, EXOC6/6B*, and *IFIH1* in the lungs of COVID-19 infected patients versus healthy controls. (B) RNA-sequence expression of *PRC1, DNER, EXOC6/6B*, IFIH1 and and KCNJ11 in nondiabetic human pancreatic islets (*n* =67) (C) Mean expression of *EXOC6* in diabetic (*n* = 12) and nondiabetic (*n* = 67) human islets.

### Gene expression of the identified shortlisted genes in the lungs of COVID-19 patients compared to healthy controls

The expression profile of the identified genes was examined in healthy versus COVID-19 infected lungs by retrieving publicly available transcriptomic data as previously described [20] (Figure3). GSE147507 dataset used for that contains RNA-sequencing of uninfected lungs compared to COVID-19 lung biopsies[21].

## Results

### SARS-CoV-2 infected hPSC-derived pancreatic organoid disturbs genes involved in insulin secretion and diabetes pathway

To understand the molecular impact of SARS-CoV-2 on genes related to the regulation of glucose hemostats, insulin secretion, or diabetes, we explored a recently uploaded transcriptomics dataset (GSE151803) of SARS-CoV-2 infected hPSC-derived pancreatic organoids. As shown in Table 1, genes related to insulin secretion [*RYR2, PCLO, PLCB4, ABCC8, KCNMA1, PRKCA, ADCY2, G*.*C*.*G*., *ATP1B2, KCNN3, ATP2B2, ATP1B2, and ATP2B1, ITPR1, ATP2B4, PRKCA*, and *RAPGEF4*] were significantly downregulated while genes related to Type I DM were upregulated [*GAD1*[22], *HLA-B*[23], *HLA-C*[23], *HLA-A*[24], *TNF*[25, 26], and *HLA-E*[27]].

**Table 1:** List of genes and enriched pathways in hPSC-derived pancreatic organoid infected with SARS-CoV-2.

| term_name | zscore | combined_score | F.D.R. | p-value | overlapping_genes | gene_set_library | log10 P | geneset |
| --- | --- | --- | --- | --- | --- | --- | --- | --- |
| Insulin secretion Homo sapiens hsa04911 | 5.176471 | 60.38258 | 0.002517 | 8.59E-06 | ['RYR2', 'PCLO', 'PLCB4', 'ABCC8', 'KCNMA1', 'PRKCA', 'ADCY2', 'GCG', 'ATP1B2', 'KCNN3', 'RAPGEF4'] | KEGG Pathways | 5.065966 | downregulated |
| Type I diabetes mellitus Homo sapiens hsa04940 | 5.581395 | 40.88509 | 0.013786 | 0.000659 | ['GAD1', 'HLA-B', 'HLA-C', 'HLA-A', 'TNF', 'HLA-E'] | KEGG Pathways | 3.181314 | upregulated |

**Table 2:** List of SNPs in *EXOC6/6B* found in the 4 Emarati patients.

|  | # locus | type | ref | length | genotype | gene | transcript | location | dbSNP |
| --- | --- | --- | --- | --- | --- | --- | --- | --- | --- |
| L1 | chr2:72692272 | INDEL | A | 2 | AAG/AAG | EXOC6B | NM_015189.2 | EXOC6B:intronic:NM_015189.2 | rs397829667: rs1032055165:<br>rs5832083: rs866386569 |
| L2 | chr2:72692272 | INDEL | A | 2 | AAG/AAG | EXOC6B | NM_015189.2 | EXOC6B:intronic:NM_015189.2 | rs397829667: rs1032055165:<br>rs5832083: rs866386569 |
| L2 | chr2:72741730 | SNV | A | 1 | A/T | EXOC6B | NM_015189.2 | EXOC6B:intronic:NM_015189.2 | rs768289243: rs746841688:<br>rs766828558: rs796091253:<br>rs752761914: rs56967666:<br>rs71884143: rs959522926:<br>rs751994268: rs777972875:<br>rs767636830: rs758383727 |
| L2 | chr2:72741734 | SNV | A | 1 | T/T | EXOC6B | NM_015189.2 | EXOC6B:intronic:NM_015189.2 | rs746841688: rs70963128:<br>rs751994268: rs796864054:<br>rs768289243: rs58454184:<br>rs71337705: rs766828558:<br>rs752761914: rs959522926:<br>rs759829795: rs777972875:<br>rs758383727 |
| O1 | chr2:72692272 | INDEL | A | 2 | AAG/AAG | EXOC6B | NM_015189.2 | EXOC6B:intronic:NM_015189.2 | rs397829667: rs1032055165:<br>rs5832083: rs866386569 |
| O2 | chr2:72411187 | SNV | T | 1 | T/C | EXOC6B | NM_015189.2 | EXOC6B:intronic:NM_015189.2 | rs141623406 |
| O2 | chr2:72723675 | SNV | A | 1 | A/G | EXOC6B | NM_015189.2 | EXOC6B:exonic:NM_015189.2 |  |
| O2 | chr2:72958333 | SNV | T | 1 | T/C | EXOC6B | NM_015189.2 | EXOC6B:exonic:NM_015189.2 | rs61736520 |

### Lung and pancreas tissues shared 540 genes with similar tissue expression levels

Next, to distinguish potential mechanisms and pathways implicated in the pathogenesis of SARS-CoV-2 infection as well as T2D, we initially explored all genes that are expressed in the lung as well as pancreatic tissues using publicly available tissue gene expression datasets (GTEx). Differential expression analysis between healthy lung samples and healthy pancreas revealed 540 genes of which their expression is not specific to lung or pancreas samples and thereby was considered as shared genes between the two tissues (Figure 1A) (Supplementary Table 1).

### Identification of SNPs associated with risk of T2D in Emirati diabetic patients

Next, using the genome-wide association studies (GWAS) database, we screened the 540 genes for SNPs that are associated with T2D risk. Only four genes (*PRC1, DNER, EXOC6/6B*, and *IFIH1*) out of the 540 shortlisted genes showed SNPs (with the lowest p-value) associated with the risk of T2D. This finding was further confirmed in the genomic profiling of the targeted next-generation sequencing from 4 Emirati T2D-patients (Figure 1B) (Supplementary Table 1). Interestingly, one insertions and deletions genetic variants (INDELs) (rs397829667:rs1032055165:rs5832083:rs866386569) in *EXOC6/6B* showed to be present in 3 of the 4 Emirati patients. Indeed, the two genes *EXOC6* and *EXCO6b*, encoded highly similar forms of the exocyst subunit *EXOC6/6b*, previously found to be essential for the exocyst-dependent exocytosis as well as insulin signaling[28].This, clearly highlighted the strong correlation between this gene and the risk of T2D.

### Expression levels of *PRC1, DNER, EXOC6/6B*, and *IFIH1* in lung tissues infected with SARS-COV-2 infection or diabetic human islets

As the data suggest that *EXOC6/6B* is a candidate link gene between T2D and COVID19, we further investigated if COVID-19 infection influences the expression of *EXOC6/6* as well as the other three shortlisted genes in lung tissues. To achieve this, we retrieved publicly available transcriptomic databases that allow us to compare healthy versus COVID-19 infected lung tissues (Figure 2A). Differential expression analysis revealed that SARS-COV-2 infection significantly reduced the expression of *PRC1, EXOC6/6B*, and *IFIH1* genes (*p* < 0.05) in the infected lung tissue compared to healthy lungs (Figure 2B). *DNER* was not present in the datasets. To gain more insights into the expression levels of the four studies genes in human pancreatic islets, we examined their expression level in healthy and diabetic islets using RNA-sequencing data (GSE50398) [18]. RNA-sequencing showed a relatively higher expression of *EXOC6/6B* in healthy/nondiabetic human islets when compared with *PRC1, DNER, IFIH1*, or the ion channel gene *KCNJ11* (a functional marker for β-cells) (Figure 2B). We did not observe any expression differences of *EXOC6/6B* between diabetic and nondiabetic islets.

## Discussion

Several reports showed an essential link between T2D and the risk of a more aggressive form of COVID-19 as well as higher mortality rates[8, 11, 29]. While many mechanisms were proposed to play a role in the increased disease severity, including dysregulated immune responses as well as abnormal inflammatory responses[8], no specific mechanisms were clearly defined to explain this clinical correlation. In addition, the effect of the virus itself on insulin secretion is also largely unknown[14]. Using a combined *in silico* and *in vivo* validation approach, we tried to identify common pathways and genes that might be involved in increasing the susceptibility to T2D, and at the same time, which are essential for SARS-CoV-2 infection progression.

Our preliminary results showed that SARS-CoV-2 was able to disturb a group of genes that are involved in the pancreatic β-cells function. This observation goes with previous reports that showed evidence that the binding of SARS coronavirus to *ACE2* receptors might lead to damage to the pancreatic islets leading to acute insulin-dependent T2D [30]. Similarly, another recent study also suggested high *ACE2* expression, which facilitates pancreatic tissue damage following SARS-CoV-2 infection[31].

Moreover, our results also showed, out of the 540 shared genes between lung and pancreas, four genes have SNPs associated with T2D in GWAS, and were confirmed in the genomic profiling of 4 Emirati patients. This includes the *EXOC6/6B* gene showed to have on indel in 3 of the 4 Emirati patients. Several reportes demonstrated a significant association between INDELs and the risk of T2D [32, 33]. INDELs have significant functional impact on gene transcription, translation efficiency, mRNA mRNA or proteins activity via changeing their structure. Phase 3 1000 Genomes Project showed that human genome has almost 3.4 million bi-allelic INDELs[34]. However, it is unclear whether INDELs are more likely to influence complex traits than SNPs. Genetic variants, such as non-coding SNPs, indels, copy number polymorphisms, or de novo mutations, may affect islet cell function and ciliated lung epithelium. Consequently, future studies such as tissue-specific knockout models is needed to determine the contribution of *EXOC6/6B*, and other genetic variations on insulin secretion and ciliated lung epithelium.

*EXOC6/6B* is part of the SEC6-SEC8 exocyst complex present in mammalian and an essential component for insulin granule docking and secretion[35-38]. Morover, *EXOC6/6B* genes, were found to encode highly similar forms of the exocyst subunit EXOC6/6b that previously identified to play a role in insulin signaling through its interaction with the glucose transporter GLUT4 [28].This further highlighted its possible role in the pathophysiology of T2D. Interestingly, some reports have also characterized *EXOC6/6B* and other members of the exocyst complex as cilia-associated proteins with ciliogenesis-associated roles[39]. Indeed, this process was also found to be essential for cystogenesis and tubulogenesis, which are essential step in the the development of many epithelial organs including lung and kidney[40]. A recent report also highlights *EXOC6/6B* as one of the top genes that might be involved in promoting the malignant transformation of human bronchial epithelial cells (HBECs)[41]. Interestingly, recent findings indicate other roles for the exocyst complex besides secretion. Among these diverse roles are epithelial polarization, morphogenesis and homeostasis. Due to its role in cell-cell interactions, it is believed that the exocyst complex may play an important role in the epithelium barrier integrity [42] which may prove to have significant implications in the lung. Another important role of the exocyst system is its role in cell migration. In a study on invasive cell lines which do not express cadherin, which is involved in cell-cell adhesion, exocyst complex relocates to protrusions that inhibit cell migration [43].It is involved in the delivery of integrins to the leading edge of migrating cells and a distrurbance in this complex leads to defects in cell motility [43-45]. Exocytosis is the mechanism used for secretion of surfactant, an important regulator of surface tension in the lungs, via fusion of surfactant-containing vesicles and the plasma membrane of alveolar type II cells. Surfactant is also proving to be involved in alveolar host defense. Therefore, any disturbance in the regulation of exocytosis may have implications on the secretion of surfactant which can lead to severe lung disorders [46]. For that reason, our finding that SARS-COV-2 infection modulates *EXOC6/6B* expression in lung tissues, might sheds light on the possible pathophysiological mechanisms shared between T2D and COVID-19.

Further investigation of those mechanisms might increase our understanding of the molecular basis behind the aggressive clinical complications observed in diabetic patients with COVID-19. Indeed, this might pave the way to discover novel therapeutic targets for a personalized and active response to the COVID19 pandemic.

## Conclusion

Our results showed that *EXOC6/6B* might be an important molecular link between insulin secretion defect and dysfunctional ciliated lung epithelium that makes diabetic patients more susceptible to SARS-COV-2 with more possible aggressive scenarios.

## Acknowledgments

We would like to thank Alagappan Kumarappan in sample processing and Dr. Rifat Hamoudi, Dr.Thenmozhi Venkatachalam, for the targeted NGS.

## Author Contributions

Conceptualization, M.Y.H., I.Y.H., S.H., and N.S.; Methodology, M.Y.H., I.Y.H., S.H.; Software, M.Y.H., I.Y.H., S.H.; Writing—Original Draft Preparation, M.Y.H., I.Y.H., S.H., J.T.; Writing—Review & Editing, M.Y.H., I.Y.H., S.H., J.T., N.S., and J.T.; Project Administration, M.Y.H., I.Y.H., S.H., J.T., N.S.; Funding Acquisition, N.S. and J.T. All authors have read and agreed to the published version of the manuscript.

## Availability of data and material

All data generated or analyzed during this study are included in this published article

